# *Pseudomonas syringae* infectivity correlates to altered transcript and metabolite levels of *Arabidopsis* Mediator mutants

**DOI:** 10.1101/2023.11.03.565469

**Authors:** Jeanette Blomberg, Viktor Tasselius, Alexander Vergara, Fazeelat Karamat, Qari Muhammad Imran, Åsa Strand, Martin Rosvall, Stefan Björklund

## Abstract

Rapid metabolic responses to pathogens are essential for plant survival and depend on numerous transcription factors. Mediator is the major transcriptional co-regulator for integration and transmission of signals from transcriptional regulators to RNA polymerase II. Using four Arabidopsis Mediator mutants, *med16*, *med18*, *med25* and *cdk8*, we studied how differences in regulation of their transcript and metabolite levels correlate to their responses to *Pseudomonas syringae* infection. We found that *med16* and *cdk8* were susceptible, while *med25* showed increased resistance. Glucosinolate, phytoalexin and carbohydrate levels were reduced already before infection in *med16* and *cdk8*, but increased in *med25*, which also displayed increased benzenoids levels. Early after infection, wild type plants showed reduced glucosinolate and nucleoside levels, but increases in amino acids, benzenoids, oxylipins and the phytoalexin Camalexin. The Mediator mutants showed altered levels of these metabolites and in regulation of genes encoding key enzymes for their metabolism. At later stage, mutants displayed defective levels of specific amino acids, carbohydrates, lipids and jasmonates which correlated to their infection response phenotypes. Our results reveal that *MED16*, *MED25* and *CDK8* are required for a proper, coordinated transcriptional response of genes which encode enzymes involved in important metabolic pathways for Arabidopsis responses to *Pseudomonas syringae* infections.

**HIGHLIGHT:** Plants need to defend themselves against different types of infections. We show that subunits of the Mediator transcriptional coactivator coordinate metabolic responses of *Arabidopsis thaliana* to infections by *Pseudomonas syringae*.

## INTRODUCTION

Mediator is an important transcriptional co-regulator complex required for integration and transmission of signals from promoter-bound transcriptional regulators to the RNA polymerase II (Pol II) transcription machinery in eukaryotic cells (Thompson et al., 1993; Kim et al., 1994). A combination of genetic, biochemical and structure biology analyses have shown that Mediator is composed of three modules; Head, Middle and Tail (Dotson et al., 2000). In addition, a fourth more loosely associated, regulatory Kinase module has been identified (Liao et al., 1995). Tail subunits are main receivers of signals from transcriptional regulators, while the Middle module transfer signals from Tail to the Head module which in turn makes direct contact with Pol II. Plant and human cells comprise more Mediator subunits than yeast, but its overall structure is conserved (Tsai et al., 2014).

Several reports have identified the plant Mediator subunits MED8, MED15, MED16, MED18, MED20, MED25 and CDK8 as important for proper immune responses to diverse types of infections (Kidd et al., 2009; Wathugala et al., 2012; Caillaud et al., 2013; Zhu et al., 2014; Fallath et al., 2017; Li et al., 2018). Evidently, each subunit has specific functions but the same subunit can have both positive and negative effects on expression of a specific gene, depending on the infection type. Most reports of Mediator subunits involvement in immune responses have focused on their transcriptional effects. Only few studies describe how Mediator mutants affect immune responses at the metabolite level.

Pathogen attack in plants is sensed by innate immune receptors present on the host cell surface or in the cytoplasm. Host surface receptor binding of microbial antigens called pathogen-associated molecular patterns (PAMPs) induces PAMP-triggered immunity (PTI) (Zhang and Zhou, 2010), while recognition of pathogen derived effectors by intracellular nucleotide-binding/leucine-rich-repeat (NLR) receptors induces an effector-triggered immune (ETI) response (Jones and Dangl, 2006). Signaling cascades from the PTI and ETI receptors lead to transcriptional reprogramming of several genes and constitute the base for a defense system in which numerous defense-related metabolites are produced.

Salicylic acid (SA) and jasmonic acid (JA) are well-studied hormones that activate distinct defense pathways. SA plays key roles in defense against hemibiotrophs and biotrophs, such as *P. Syringae*, whereas JA acts together with ethylene to induce resistance against necrotrophic pathogens. These pathways work in parallel by negatively regulating each other. Biosynthesis of SA during infection is mainly promoted by induced expression of *ISOCHORISMATE SYNTHASE1 (ICS1)*, which in turn is positively regulated by *ENHANCED DISEASE SENSITIVE 1 (EDS1)* and *PHYTOALEXIN-DEFICIENT4 (PAD4)*. Signaling downstream from SA is executed by activation of *NONEXPRESSOR OF PATHOGENESIS-RELATED GENES1 (NPR1)* which translocates from the cytoplasm to the nucleus to induce expression of various defense genes, for example *NIMIN1* and pathogen regulated (PR) genes. Reprogramed transcription of defense genes induces a long-term memory which is distributed throughout the entire plant, termed systemic induced response (SAR). Thus, production of SA and signaling from SA is critical for functional PTI, ETI, and SAR responses (Durrant and Dong, 2004).

In addition to hormone activation, plants alter their production of numerous secondary metabolites in response to pathogen attack. Glucosinolates (GLSs) constitute a large family of sulfur-and nitrogen-containing β-D-thioglucoside-*N*-hydroxysulfate defense compounds. Three GLS classes are present in Arabidopsis: methionine-derived aliphatic GLSs, phenylalanine-derived benzenic GLSs, and tryptophan-derived indolic GLSs. The most abundant foliar GLSs are derived from methionine or tryptophan by chain elongation, core structure formation and secondary modifications. These GLSs are biologically inactive but are rapidly hydrolyzed and liberated from the glucose moiety by myrosinases upon plant injury. This transforms them into active compounds, such as isothiocyanates (ITCs), thiocyanates, simple nitriles and epithionitriles. Transcription Factors (TFs) of the MYB R2R3-family have been shown to control GLS biosynthesis by activating transcription of genes encoding key enzymes in these pathways. MYB28, MYB29 and MYB76 regulate expression of *MAM1*, *CYP79F1* and *CYPF2* which encode key enzymes in the aliphatic pathways (Sønderby et al., 2010), while MYB34, MYB51 and MYB122 regulates *CYP79B2* and *CYP79B3*, which encode key enzymes in the indolic pathway. Besides indolic GLSs, other indolic phytoalexins that contribute to the defense response, like Camalexin, indole-3-carbaldehyde (ICHO), indole carboxylic acid (ICOOH) and ascorbigen, are also produced from tryptophan (Böttcher et al., 2014).

The phenylpropanoid pathway originates from phenylalanine and leads to production of secondary metabolites such as flavonoids (flavonols/kaempferols and anthocyanins), coumarins, lignans and lignin (Fraser and Chapple, 2011). Glucosyltransferases play key roles in formation of sinapates and sinapate ester intermediates, possibly through production of the high energy compound 1-O-sinapoyl glucose. It is metabolized to sinapoylmalate and sinapoylcholine which constitute the syringyl units found in lignins. Sinapate ester levels have been shown to change after infection but their relevance for pathogenicity has not yet been established.

Lipids also influence plant-pathogen interactions at various levels ranging from interaction with virulence factors to activation and control of host plant immune defenses. These include fatty acids, oxylipins, phospholipids, glycolipids, glycerolipids, sphingolipids, and sterols. Oxylipins are derived from polyunsaturated fatty acids and can be formed enzymatically or non-enzymatically. The enzymatic pathway starts with lipoxygenases and leads to formation of 12-oxophytodienoic acid (OPDA), jasmonates, aldehydes and other oxylipin metabolites such as monogalactosyldiacylglycerols (MGDGs) and digalactosyldiacylglycerols (DGDGs). Several complex lipid molecular species where fatty acids are linked to an OPDA or dinor-OPDA (dnOPDA) have been characterized (Stelmach et al., 2001; Hisamatsu et al., 2005; Kourtchenko et al., 2007).

Here we combine metabolic profiling and RNA-sequencing (RNA-seq) to reveal mechanisms for transcriptional and metabolite responses to infection caused by *P. syringae* in wild type Arabidopsis and in plants carrying mutations in Mediator subunits.

## MATERIALS AND METHODS

### Plant material and growth conditions

Wild type Columbia-0 (Col-0) *Arabidopsis thaliana* plants were used as reference genotype for susceptibly to virulent *P. syringae* pv. *tomato* DC3000 (*Pst*. DC3000). Mediator mutants in the Col-0 background: *med16* (alias sfr6-2; SALK_048091), *med18* (SALK_027178), *med25* (SALK_129555) and *cdk8* (GABI_564F11), have been described previously (Ng et al., 2013; Davoine et al., 2017; Crawford et al., 2020). Plants were grown in soil under controlled short day (8 h light /16 h dark) conditions at 22 °C and 67 % humidity.

### Bacterial infection and colony forming unit (CFU) assay

Single colonies were picked and grown in Kings Broth (KB) liquid media containing 50 µg/ml rifampicin at 28 °C over night. Cultures were diluted to OD_600_ of 0.3 and grown to an OD_600_ of 0.8-1. Exponentially growing bacteria was collected, washed twice in 10 mM MgCl_2_ and resuspended in the same solution at a concentration of 2 × 10^6^ CFU/ml. 4-6 lower leaves of 5–6-week-old plants were injected on their abaxial side with either *Pst.* DC3000 suspension (infections) or 10 mM MgCl_2_ (controls). Leaves from 2-3 plants were pooled for one biological replicate. For quantification of bacterial growth, 40 mg pooled leaf samples were grinded in 1 mL 10 mM MgCl_2_, serially diluted and plated on KB plates containing 50 µg/ml rifampicin. Bacterial numbers were determined using the following formula: CFU = ((Number of colonies x volume x dilution factor) / volume plated)/sample weight.

### Metabolite extraction

Samples were prepared from control and infected 5-week-old plants (Gullberg et al., 2004). 20 mg of grinded samples were mixed with 1 mL extraction buffer (20/20/60 v/v/v chloroform:water:methanol) including internal standards for GC-MS and LC-MS. LC-MS internal standards were: 13C9-phenylalanine, 13C3-caffeine, D4-cholic acid, D8-arachidonic acid and 13C9-caffeic acid (Sigma, St. Louis, MO, USA). GC-MS internal standards were: L-proline-13C5, alpha-ketoglutarate-13C4, myristic acid-13C3, cholesterol-D7 (Cambridge Isotope Laboratories, Inc., Andover, MA, USA) and succinic acid-D4, salicylic acid-D6, L-glutamic acid-13C5,15N, putrescine-D4, hexadecenoic acid-13C4, D-glucose-13C6, D-sucrose-13C12 from Sigma. The samples were bead-beated and centrifuged as described (Davoine et al., 2017). Most of the supernatants, 200 µL for LC-MS analysis and 50 µL for GC-MS analysis, were transferred to micro vials, evaporated to dryness and stored at -80 °C until analysis. Small aliquots of the remaining supernatants were pooled and used as quality control (QC) samples. MSMS analysis (LC-MS) was run on the QC samples for identification purposes. The samples were analyzed in batches according to a randomized run order on both GC-MS and LC-MS.

### GC-MS profiling

Derivatization and GC-MS analysis were performed as described (Gullberg et al., 2004). 0.5 μL of the derivatized sample was injected in splitless mode by an L-PAL3 autosampler (CTC Analytics AG, Switzerland) into an Agilent 7890B gas chromatograph equipped with a 10 m x 0.18 mm fused silica capillary column with a chemically bonded 0.18 μm Rxi-5 Sil MS stationary phase (Restek Corporation, U.S.) The injector temperature was 270 °C, the purge flow rate was 20 mL/min, and the purge was turned on after 60 seconds. The gas flow rate through the column was 1 mL/min. The column temperature was held at 70 °C for 2 minutes, then increased by 40 °C/min to 320 °C and held there for 2 minutes. The column effluent was introduced into the ion source of a Pegasus BT time-of-flight (TOF) mass spectrometer, GC/TOFMS (Leco Corp., St Joseph, MI, USA). The transfer line and the ion source temperatures were 250 °C and 200 °C, respectively. Ions were generated by a 70-eV electron beam at an ionization current of 2.0 mA, and 30 spectra/second were recorded in the mass range m/z 50 – 800. The acceleration voltage was turned on after a solvent delay of 150 seconds. The detector voltage was 1800-2300 V.

### LC-MS profiling

The samples were reconstituted in 20 µL of 50% (v/v) methanol before analysis. Each batch of samples were initially analyzed in positive mode, followed by a switch to negative mode for a second injection of each sample after analyzing all samples within the batch. The chromatographic separation was performed on an Agilent 1290 Infinity UHPLC-system (Agilent Technologies, Waldbronn, Germany). Two μL of each sample were injected onto an Acquity UPLC HSS T3, 2.1 x 50 mm, 1.8 μm C18 column in combination with a 2.1 mm x 5 mm, 1.8 μm VanGuard precolumn (Waters Corporation, Milford, MA, USA) held at 40 °C. The gradient elution buffers were A (0.1 % formic acid) and B (75/25 (vol/vol) acetonitrile:2-propanol, 0.1 % formic acid), and the flowrate was 0.5 mL/min. The compounds were eluted with a linear gradient consisting of 0.1 – 10 % B over 2 minutes. B was then increased to 99 % over 5 minutes and held at 99 % for 2 minutes. B was then decreased to 0.1 % over 0.3 minutes and the flow-rate was increased to 0.8 mL/min for 30 seconds. These conditions were kept for 0.9 minutes, after which the flow-rate was reduced to 0.5 mL min^-1^ for 0.1 minutes before the next injection. The compounds were detected using an Agilent 6546 Q-TOF mass spectrometer equipped with a jet stream electrospray ion source operating in positive or negative ion mode. The settings were kept identical between the modes, with exception of the capillary voltage. A reference interface was connected for accurate mass measurements. The reference ions purine (4 μM) and HP-0921 (Hexakis(1H, 1H, 3H-tetrafluoropropoxy)-phosphazine) (1 μM) were infused directly into the MS (flow rate of 0.05 mL/min) for internal calibration, and the monitored ions were purine m/z 121.05 and m/z 119.03632; HP-0921 m/z 922.0098 and m/z 966.000725 for positive and negative mode respectively. The gas temperature was set to 150 °C, the drying gas flow to 8 L/min and the nebulizer pressure to 35 PSI. The sheath gas temperature was set to 350 °C and the sheath gas flow to 11 L/min. The capillary voltage was set to 4,000 V in positive ion mode, and to 4,000 V in negative ion mode. The nozzle voltage was 300 V. The fragmentor voltage was 120 V, the skimmer 65 V and the OCT 1 RF Vpp 750 V. The collision energy was set to 0 V. The m/z range was 70 – 1700, and data was collected in centroid mode with an acquisition rate of four scans/second (1977 transients/spectrum).

### Metabolite data analysis

For the GC-MS data, all non-processed MS-files from the metabolic analysis were exported from the ChromaTOF software in NetCDF format to MATLAB R2021a (MathWorks, Natick, MA, USA), where all data pre-treatment procedures, such as base-line correction, chromatogram alignment, data compression and Multivariate Curve Resolution were performed. The extracted mass spectra were identified by comparisons of their retention index and mass spectra with libraries of retention time indices and mass spectra (Davoine et al., 2017). Mass spectra and retention index comparison was performed using the NIST MS 2.2 software. Annotations of mass spectra were based on reverse and forward searches in the library. Masses and ratios between masses indicative of a derivatized metabolite were especially notified. The mass spectrum with the highest probability indicative of a metabolite and the retention index between the sample and library for the suggested metabolite was ± 5 (usually less than 3) and the deconvoluted “peak” was annotated as an identification of a metabolite. For the LC-MS data, all data processing was performed using the Agilent MassHunter Profinder version B.10.00 (Agilent Technologies Inc., Santa Clara, CA, USA). The processing was performed both in a targeted and an untargeted fashion. For target processing, a pre-defined list of metabolites was analyzed using the Batch Targeted feature extraction in MassHunter Profinder. An in-house LC-MS library built up by authentic standards run on the same system with the same chromatographic and mass-spec settings, was used for the targeted processing (Davoine et al., 2017). The identification of the metabolites was based on MS, MSMS and retention time information. Differences in metabolite concentrations were tested using pairwise t-test assuming equal variance between groups. P-values below 0.05 were considered significant and due to the exploratory nature of the metabolite analysis no correction for multiple testing was performed.

### RNA isolation and qPCR

Total RNA was extracted from 100 mg of grounded leaf tissue using the E.Z.N.A Plant Mini Kit (Omega Bio-tek, Norcross, USA) and contaminating DNA was removed using turbo DNAfree DNAse I (Ambion, Foster City, USA). Total RNA (1 µg) was reverse transcribed using iScript reverse transcription supermix (Biorad, Solna, Sweden). RT-qPCR was performed using a LightCycler 96 and the PowerUp SYBR green master mix (Applied Biosystems, Massachusetts, USA). Gene expression levels were normalized to the reference genes *RCE1* (AT4G36800) and *ACT2* (AT3G18780) and displayed as relative units. Three biological and two technical replicates were used for each sample. RT-qPCR sequence primers are shown in Supplementary Table S1.

### RNA-sequencing

RNA-seq data for uninfected Col-0, *med16*, *med18* and *cdk8* plants have been published (Crawford et al., 2020). RNA-seq data for uninfected *med25* and Col-0 were obtained in an equivalent manner. In brief, isolated and DNAse treated RNA extracted from 5-week-old plants was assayed for RNA integrity with the Agilent 2100 Bioanalyzer using the RNA Nano 6000 kit (Agilent Technologies, Santa Clara, USA). Single-end RNA-seq was performed on a HiSeq 2500 High Output V4 platform (Illumina, San Diego, USA), generating 13–32 million reads.

### Analysis of transcriptomic data

The raw RNA-seq data was pre-processed by NGI Uppsala using TrimGalore and FastQC. Reads were aligned to the Araport11 reference genome and read counts were obtained using the kallisto R package (v0.45.1) (Bray et al., 2016). To analyze differential gene expression, the R DESeq2 package (v1.30.0) (Love et al., 2014) was employed. Using pairwise differential expression analysis comparisons, fold changes were obtained between the mutant genotypes and their respective wild-type controls (Supplementary Table S2). Thus, a comparison was made between mutants and wild-type plants under normal, non-stress conditions.

### Gene Ontology (GO) functional enrichment analyses

GO functional enrichment analyses was done using Genomes (KEGG) pathway enrichment (www.ncifcrf.gov), with the whole genome background and a cutoff of an adjusted p-value of 0.05 was applied to account for multiple hypothesis testing. Lists of enriched GO functional categories were reduced by removal of redundant categories using REVIGO (http://revigo.irb.hr) with default settings and the database set to “Arabidopsis thaliana”. Overlaps between gene sets were determined using http://bioinformatics.psb.ugent.be /webtools/Venn.

## RESULTS

### *med16* and *cdk8* show increased, and *med25* reduced, susceptibility to infection by *P. syringae*

Arabidopsis *med16*, *med25*, *med18*, and *cdk8* mutants, representing each of the four tail, middle, head, and kinase Mediator modules, display diverse defects in defense signaling in response to different types of infection (Kidd et al., 2009; Chen et al., 2012; Wathugala et al., 2012; Lai et al., 2014; Zhu et al., 2014). To investigate their responses to *P. syringae*, we infected Col-0 and each mutant with *P. syringae* pv. *tomato* DC3000, a virulent strain that suppresses PTI by injecting effectors into Arabidopsis cells via bacterial secretion systems. This generates a mild disease response that becomes visible 2-3 days after infection in the form of wet, chlorotic, and spreading necrotic lesions (Xin and He, 2013). The mutants also display flowering-time phenotypes (Cerdán and Chory, 2003; Knight et al., 2008; Zheng et al., 2013). To avoid indirect developmental effects, we infected our plants under non-inductive, short-day conditions at the age of 5-6 weeks (mature rosette stage). The number of CFUs for each line was determined 72 hours post infection (p.i.) using serial dilution (Fig. 1A-1B). *med16* and *cdk8* showed significantly higher, and *med25* lower, CFUs relative to Col-0. A slight increase of CFUs was detected also in *med18* but in accordance with other studies, this increase was not statistically significant (Zhang et al., 2012). The susceptibility of *med16* and *cdk8* was also detected as more severe disease symptoms of leaves at 72 hours p.i. (Fig. 1C).

**Figure 1.**
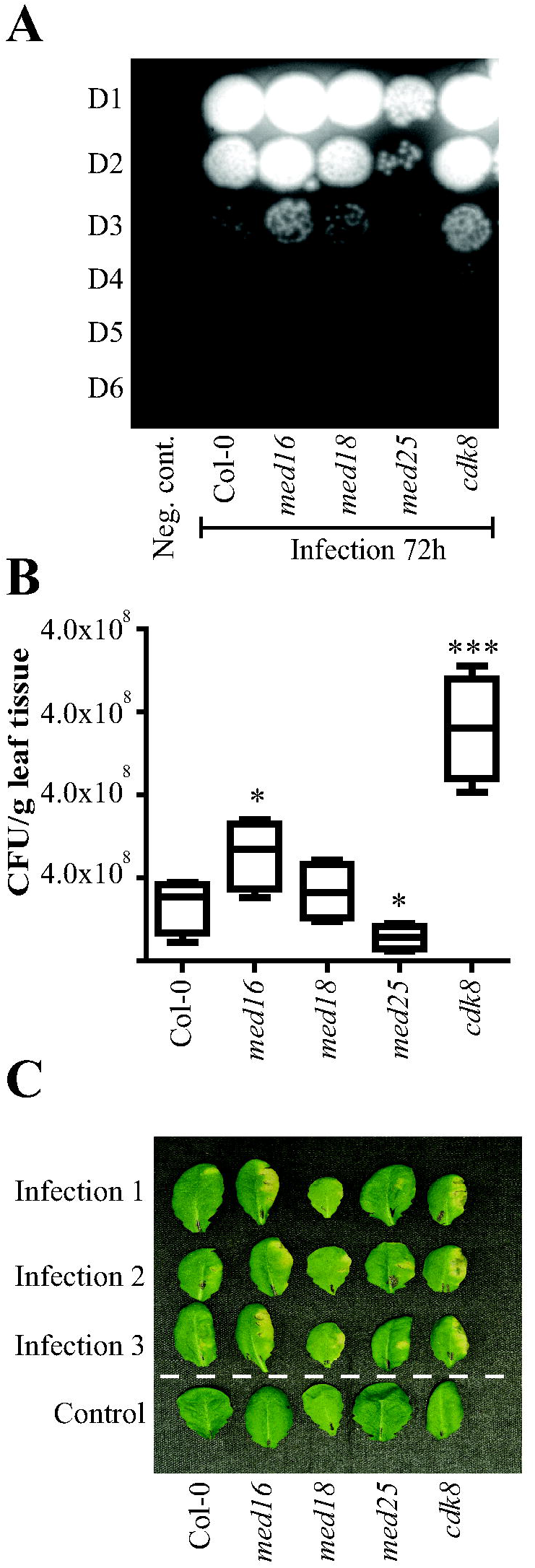
Susceptibility of Col-0 and Mediator mutants to infection by *P. syringae*. Four leaves each of 5-week-old plants were injected with either 10 mM MgCl_2_ (control) or 1 x 10^6^ CFU/ml of the virulent *Pst.* DC3000. (**A**) Bacteria were extracted from 100 mg of grounded tissue from mock-infected and infected plants and colonies formed from 10 x serial dilutions (D1 to D6) at the 72-hour time point are shown (**B**) The box plot represents the means of CFUs of the D3 dilution from four independent infections. Asterisks indicate significant differences between mutants and Col-0 (Student’s t-test, *: p< 0.05, **: p< 0.01, ***: p< 0.001) and the error bars show mean standard deviation (SD) of four biological replicates. (**C**) An illustrative picture of disease symptoms visible as necrotic and chlorotic lesions on leaf surfaces from three independently infected plants (infection 1-3) and one control experiment.

### Global metabolite levels in Col-0 and Mediator mutants deviate between infected and mock-infected leaves with the highest differences detected 72 hours after infection

Untargeted LC-MS and GC-MS were performed at 24 and 72 hours p.i. to determine global differences in metabolite levels between Col-0 and mutants in both infected and mock-infected lines. We recorded 171 and 75 metabolites from the LC-MS and GC-MS, respectively (Supplementary Table S3). For twenty-four metabolites identified in both assays, we decided to use the LC-MS data resulting in a list of totally 222 metabolites for further analyses.

An overview of metabolite levels in Col-0 and mutants at 24 and 72 hours after infection or mock-infection is illustrated as a principal component analysis (PCA) in Fig. 2A. PC1 and PC2 accounted for 42.08% and 11.41% of the total variation, respectively. The metabolite profiles in mutants differed slightly from Col-0 under control conditions (PC2) while major differences were detected after infection, especially at 72 hours p.i. (PC1). The four biological replicates for each mutant and treatment grouped well together, indicating high reproducibility.

**Figure 2.**
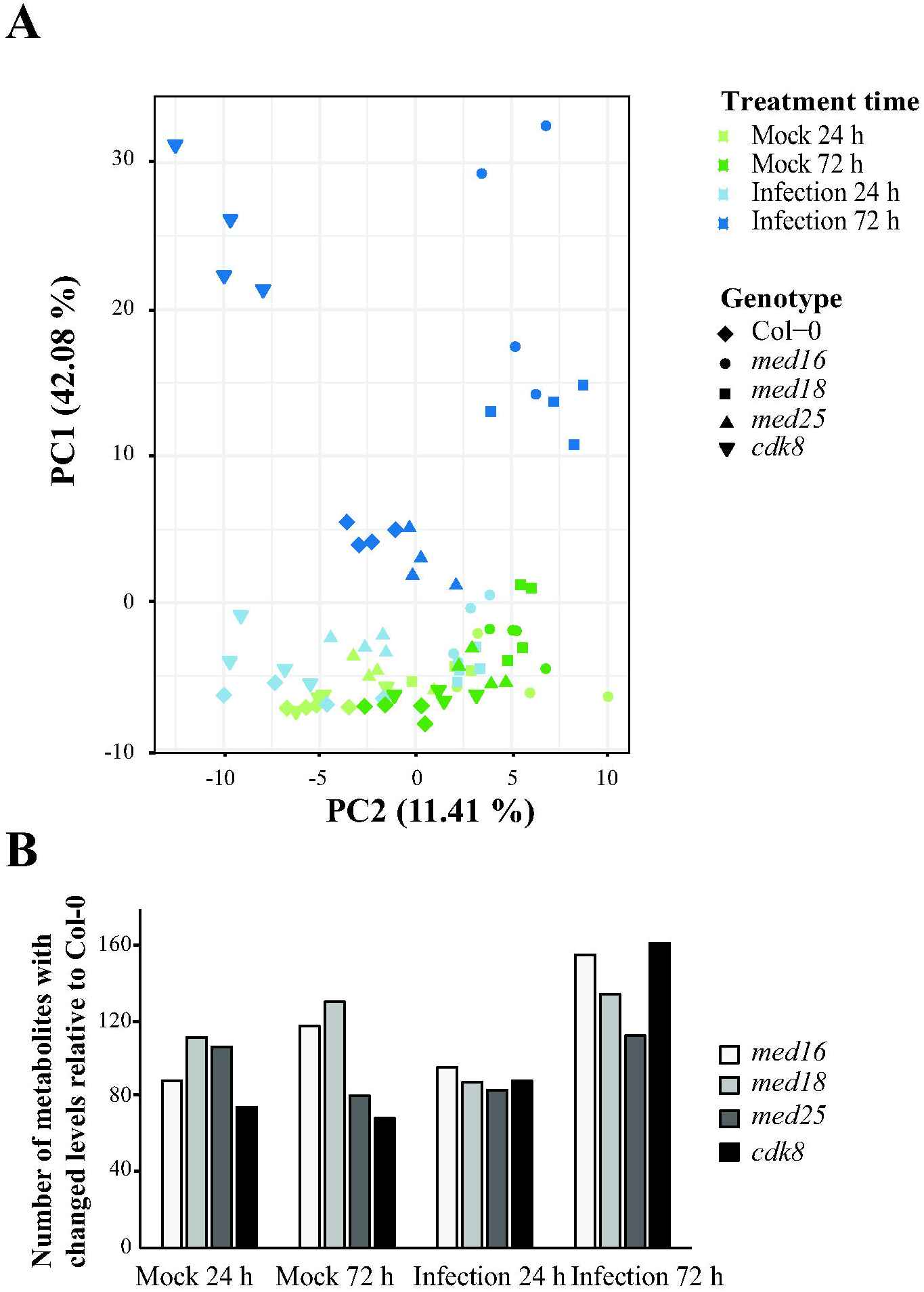
Non-targeted metabolomic profiling of Col-0 and Mediator mutants at the 24-and 72-hour time points after infection. (**A**) Multivariate statistical analysis of GC-MS and LC-MS non-targeted metabolite profiles of infected and mock-infected Col-0, *med16*, *med18*, *med25* and *cdk8* leaves at the 24-and 72-hour time points are shown as a PCA score plot. Each data point represents the entire metabolome (GC-MS plus LC-MS) for each replicate and are arranged in the first and second dimensions (x and y respectively). (**B**) Number of metabolites with statistically significant (Student’s t-test, p< 0.05) altered levels in mutants compared to Col-0 at the indicated time points and treatments.

In agreement with their increased susceptibility, *med16* and *cdk8* showed the largest number of changed metabolite levels compared to Col-0 at both 24 and 72 hours p.i. (Fig. 2B). Lists of the normalized peak values of all metabolites in infected and mock-infected Col-0 and mutants at each time point are shown in Supplementary Table S3. Ratios between metabolite levels in each mutant at each time point relative to Col-0 are shown in Supplementary Table S4. Differences in metabolite levels were further visualized using heatmaps and hierarchical clustering (Supplementary Fig. S1). The heatmap confirms that the more sensitive mutants, *med16* and *cdk8*, in general display a pronounced increase in levels of secondary metabolites at 72 hours p.i., in particular of sugars, lipids, and amino acids.

### Uninfected Col-0 and Mediator mutants display differences in metabolite levels in specific pathways

To identify metabolites that differ between mock-infected Col-0 and each mutant, those that displayed a significant difference (log_2_-fold change (FC) > 0.5 or <-0.5; p-value < 0.05) at both time points were identified. *med16* showed the highest number of decreased metabolites (thirty-two) and *med18* the highest number of increased metabolites (thirty-nine) (Fig. 3A). To comprehensively illustrate these differences, we grouped the 222 metabolites into nineteen categories according to their classification in the Human Metabolome Database (HMDB; https://hmdb.ca/) (Supplementary Table S5). The mutants displayed significant differences in levels of metabolite categories relative to Col-0 already in mock-infected lines (Fig. 3B). In general, phytoalexins, carbohydrates and GLSs were reduced in the susceptible *med16* and *cdk8*, while they were increased in *med18* and/or the more resistant *med25*. In support of these metabolite assays, RNA-seq analyses showed that *med16* and *cdk8* displayed a more pronounced reduction of transcripts encoding enzymes in GLS biosynthesis (Fig. 3C). We also noticed reduction in *med16* and *cdk8* of transcripts belonging the GO categories “Glycine, serine and threonine metabolism”, “Sulfur metabolism” and “2-oxocarboxylic acid metabolism”, which represent pathways linked to GLS metabolism. Finally, *med16*, *med18* and *cdk8* showed defects in expression of genes in the category “plant-pathogen interactions”, albeit in different directions where *med18* and *cdk8* showed decreased, while *med16* showed increased levels.

**Figure 3.**
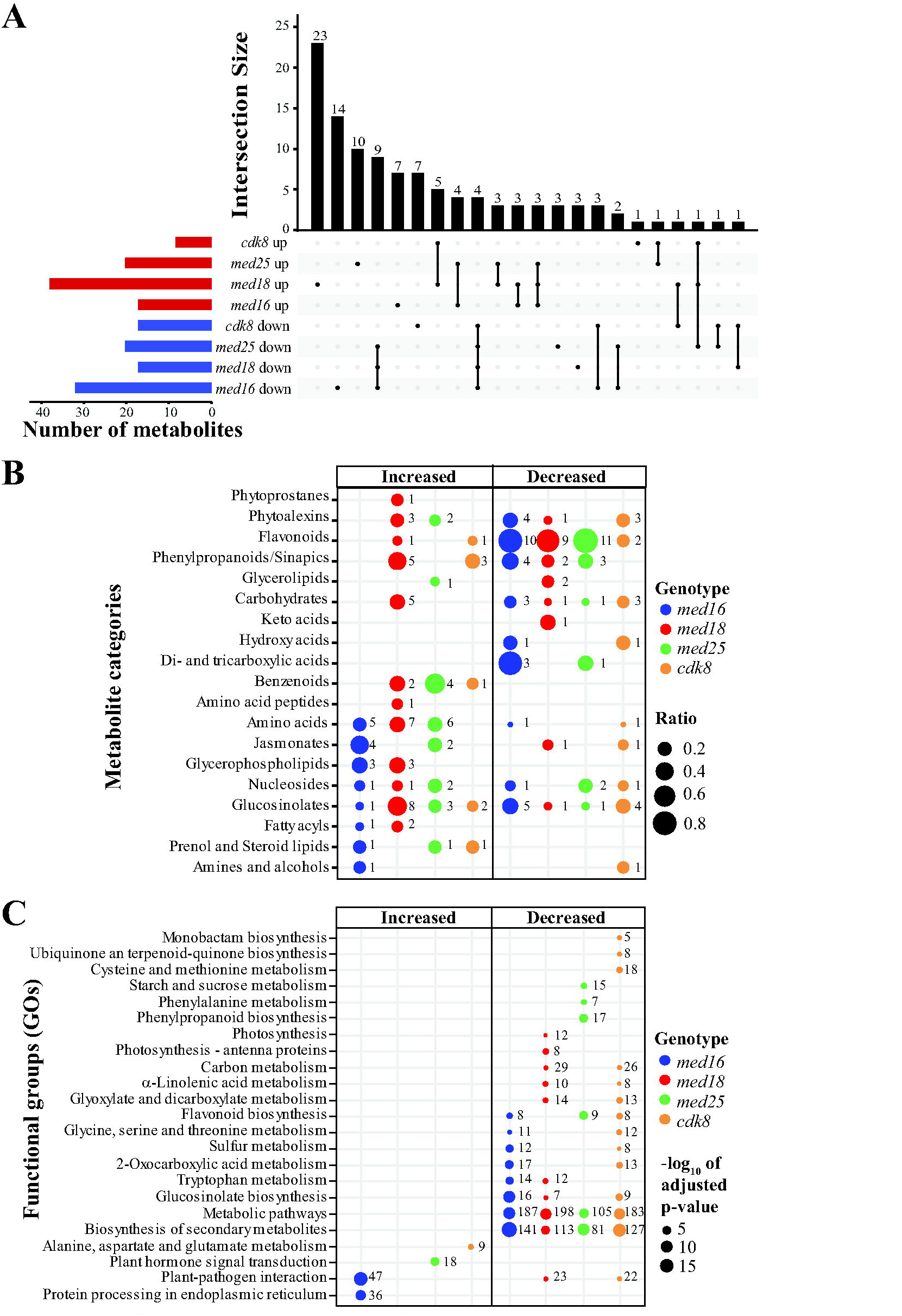
Comparison of metabolites and mRNAs with altered levels in Mediator mutants relative to Col-0 in mock-infected plants. (**A**) Upset diagram illustrating both the overlap and uniqueness of metabolites that display altered levels in mock-infected mutants relative to Col-0. Only metabolites that show statistically significant differences (log_2_-FC > 0.5 or <-0.5 and p < 0.05) at both the 24-and 72-hour time points are included. The total number of metabolites that display altered levels in each mutant are represented as red (increased) and blue (decreased) bars to the left. (**B**) Metabolites that displayed statistically distinct levels in mock-infected mutants relative to mock-infected Col-0 were grouped into nineteen metabolite categories (see Supplementary Table S5). Sizes of circles represent the ratio of the of the number of metabolites in each category that display significantly different levels in mutants relative to Col-0 and the total amount of metabolites detected for each category in our full data set of 222 metabolites. Numbers to the right of each circle represent the total number of metabolites that show significantly changed levels in each mutant. (**C**) GO enrichment analysis of alterations in defined KEGG-pathways of differentially expressed genes (DEGs) in each mutant relative to Col-0. The sizes of circles represent the significance (-log_10_ of Benjamini-Hochberg adjusted p-value) of the enrichment for each GO and numbers to the right of circles represent the number of DEGs for each mutant in the respective GO.

As shown above, *med16* and *cdk8* are susceptible, while *med25* shows increased resistance to *P. syringae* infection relative to Col-0. We therefore focused on metabolites that display similar differences in *med16* and *cdk8* relative to Col-0, already in mock-infected plants, and those that where uniquely different from mock-infected Col-0 in each of *med16*, *med25* and *cdk8* at both timepoints. No metabolites were commonly increased in *med16* and *cdk8* relative to Col-0, but two GLSs (Sulforaphane and a Sulforaphane fragment) and one phytoalexin (Benzyl dithiocarbamate) were commonly decreased and might contribute to their susceptibility (Supplementary Table S6). To get a broader view of GLSs levels in mutants, we calculated their mean values in mock-infected mutants relative to Col-0 at the 24-and 72-hour time points. Five GLSs (Sulforaphane, a Sulforaphane fragment, Hirsutin, a Hirsutin fragment and Neoglucobrassicin) displayed reduced levels in the sensitive *med16* and *cdk8* (Supplementary Fig. S2A). In contrast, GLS levels were increased in *med18* and *med25*. We also detected several phytoalexins that showed reduced levels in *med16* and *cdk8* but increased in *med18* and *med25* (Supplementary Fig. S2B).

Fourteen metabolites were uniquely decreased in mock-infected *med16* and eight in *cdk8* relative to Col-0 at both time points (Supplementary Table S6). They represent various metabolite categories, but we noticed that five (*med16*) and three (*cdk8*) were GLSs or phytoalexins. By instead using mean values of levels at 24-and 72-hours p.i., we identified five (*med16*) and six (*cdk8*) downregulated GLSs, and four (*med16*) and three (*cdk8*) downregulated phytoalexins (Supplementary Fig. S2A-S2B). In line with the opposite phenotype of *med25* relative to *med16* and *cdk8*, GLSs and phytoalexins were typically upregulated in *med25*. None of the metabolites that were uniquely decreased *med25*, and none of the uniquely increased metabolites in *med16* and *cdk8* were GLSs or phytoalexins (Supplementary Table S6). Thus, GLS and phytoalexin levels in mock-infected mutants correlate with their susceptibility to *P. syringae* infection. Finally, ten metabolites were uniquely increased in *med25*. The most common category was benzenoids (4 metabolites; Supplementary Fig. S2C) which are derivatives of SA.

### Reduction of metabolite levels during the early infection response in Col-0 is impaired in *med16* and *med18*

Hierarchical clustering of all metabolites that showed changed levels in Col-0 at 24 hours p.i. relative to its control, revealed a set of decreased GLSs and nucleosides whereas phytoprostanes, lipids, amino acids, benzenoids, jasmonates, and phytoalexins were increased (Fig. 4A). Mock-infected plants were used as controls rather than untreated, since the mechanical manipulation connected with the infection procedure could elicit an unwanted wounding response. The reduction of GLSs detected in Col-0 was completely absent in *med16* and impaired in *med18*. Similarly, the reduction in nucleosides levels observed in the early infection response in Col-0, was absent in *med16* and defect in the other mutants (Supplementary Table S7).

**Figure 4.**
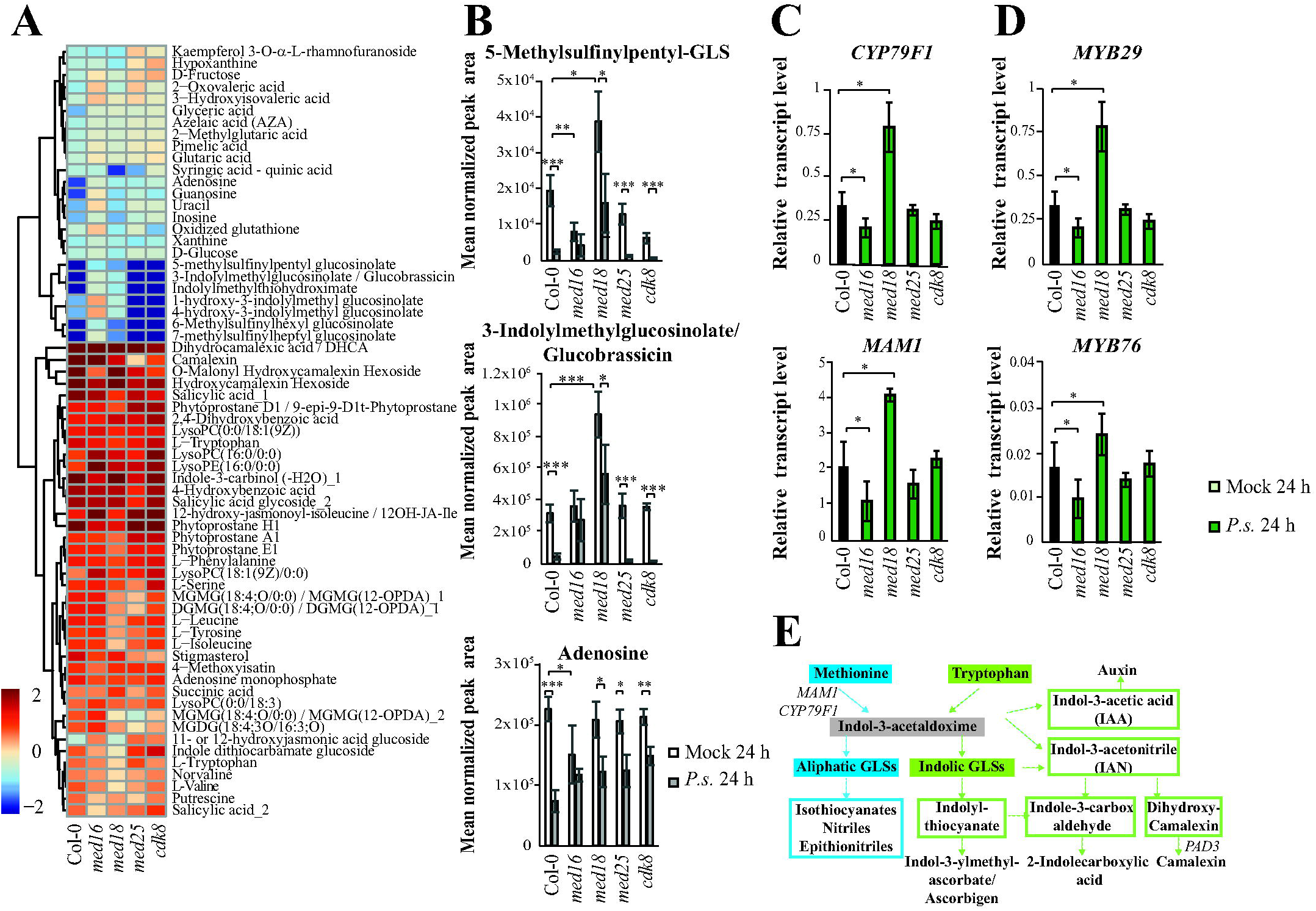
Impaired reduction of specific metabolite levels is caused by different mechanisms in *med16* and *med18*. (**A**) Heat map and hierarchical clustering of metabolites that show a log_2_-FC >0.5 or <-0.5, and p < 0.05 in infected relative to mock-infected Col-0 at the 24-hour time point. (**B**) Bar graphs of normalized mean peak areas from four biological replicates of infected and mock-infected Col-0 and mutants at the 24-hour time point. (**C**) Quantification of transcript levels for two of the aliphatic-GSL biosynthesis enzymes (*CYP79F1* and *MAM1*) in uninfected Col-0 and mutants using RT-qPCR. (**D**) Quantification of transcript levels for two of the aliphatic-GSL biosynthesis regulating transcription factors (*MYB29* and *MYB76*) in uninfected Col-0 and mutants using RT-qPCR. Asterisks indicate significant differences (*: p< 0.05, **: p< 0.01, ***: p< 0.001) and error bars represent the SD of four biological replicates for metabolite levels and three biological replicates for transcript levels. **E**) Simplified overview of the aliphatic-GLS, indolic-GLS and phytoalexin synthetic pathways. Each arrow indicates multiple enzymatic steps and only the enzymes analyzed here are indicated.

Lack of metabolite reduction in mutants can result from two different mechanisms. One possibility is that a metabolite level is reduced already in untreated mutants relative to Col-0, and remains at this low level after infection. Alternatively, mutants could have normal levels of the metabolite before infection but are unable to reduce the levels in response to infection. To distinguish between these possibilities, we compared the absolute levels for one representative metabolite each of the two GLS classes: 5-methylsulfinylpentyl-GLS (aliphatic) and 3-Indolylmethylglucosinolate/Glucobrassicin (indolic), as well as for one nucleoside (adenosine). We found that the mechanisms for the lack of metabolite reduction in response to infection differed between mutants and metabolites (Fig. 4B). *med16* showed lack of reduction of aliphatic GLSs and nucleosides at 24 hours p.i. due to reduced levels already in the mock-infected controls. In contrast, *med16* was deficient in reduction of Glucobrassicin in response to infection. Mock-infected *med18* exhibited elevated levels of both aliphatic and indolic GLSs, as we have reported previously (Davoine et al., 2017). This results in inadequate down-regulation of these metabolites in *med18* which after infection displayed GLSs levels similar to those observed in mock-infected Col-0 (Fig. 4B). Metabolites showing a log_2_FC <-1 in Col-0, and their fold changes in each mutant, are shown in Supplementary Table S7. For the full set of log_2_FCs of all 222 metabolites in all mutants relative to Col-0, see Supplementary Table S4.

We used qPCR to quantify mRNA levels encoding key enzymes in the aliphatic GLS pathway in mock-infected Col-0 and mutants (Fig. 4C). *CYP79F1* and *MAM1* levels were increased in *med18* but decreased in *med16* which correlated with their aliphatic GLS levels (Fig. 4B). MYB28, MYB29 and MYB76 belong to the R2R3-family of MYB TFs which regulate expression of *CYP79F1* and *MAM1* (Sønderby et al., 2010). In line with the effects observed for their target genes, the *MYB29* and *MYB76* mRNA levels were decreased in mock-infected *med16* but increased in *med18* relative to Col-0 (Fig. 4D). For an overview of the GLS and phytoalexin metabolic pathways, see Fig. 4E.

### Metabolites induced in Col-0 during the early response show diverse types of impaired induction in mutants

Representative metabolites that were increased at 24 hours p.i. in Col-0 but showed no response in mutants are shown in Fig. 5. All significantly increased metabolites in Col-0 at 24 hours p.i. relative to control, and their levels in each mutant are shown in the heat map in Fig. 4A. Metabolites that showed a log_2_FC >1 in Col-0, and their fold changes in each mutant are listed in Supplementary Table S8.

**Figure 5.**
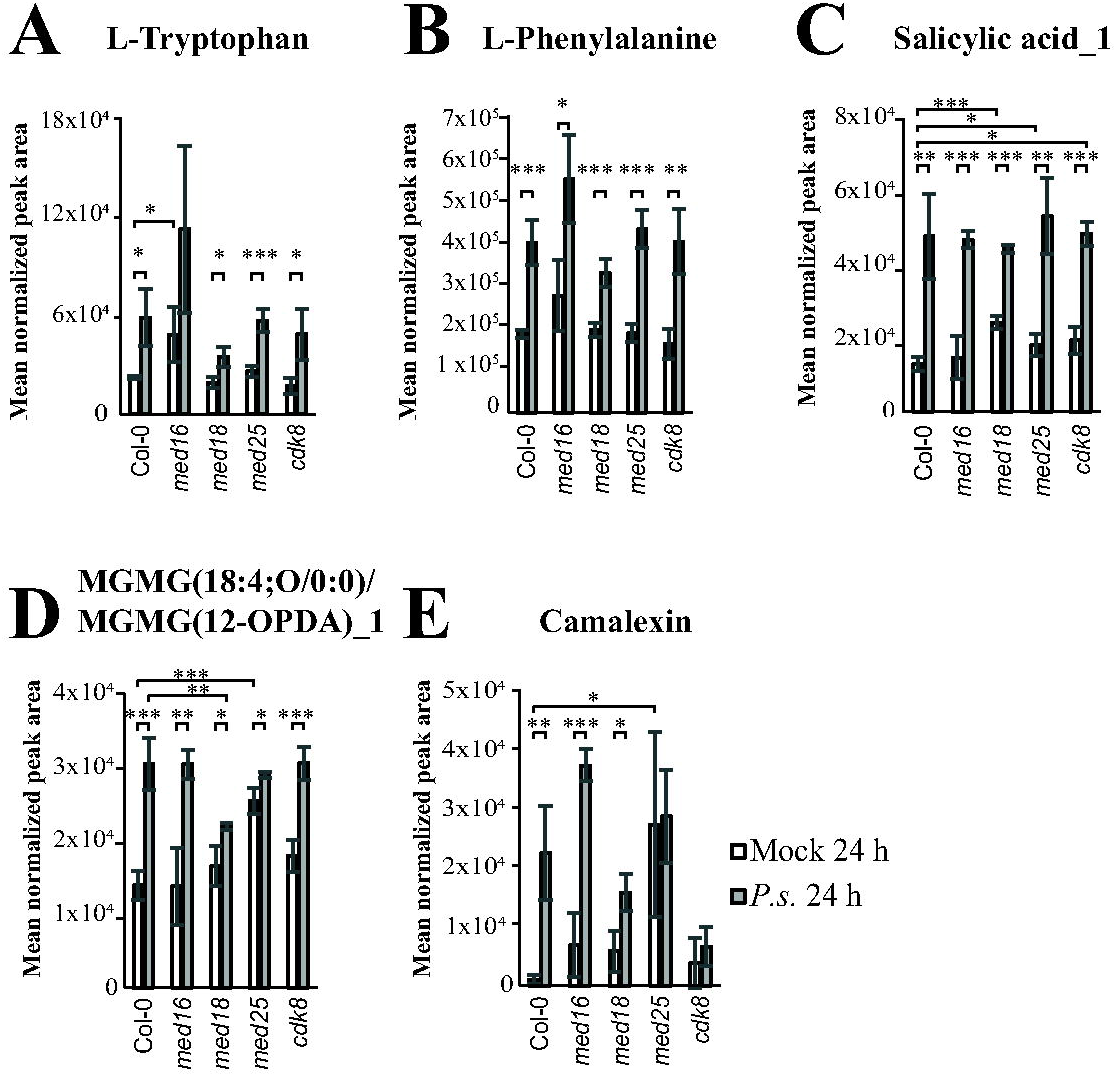
All mediator mutants have unique alterations in the induction of specific metabolites compared to Col-0 at the early time point after infection. Metabolite levels of L-Tryptophan (**A**), L-Phenylalanine (**B**), Salicylic acid_1 (**C**), MGMG (18:4;O/0:0) (**D**), and Camalexin (**E**) in infected and mock-infected Col-0 and mutants at the 24-hour time point. Asterisks indicate significant differences (*: p< 0.05, **: p< 0.01, ***: p< 0.001) and the error bars show the SD of four biological replicates.

A set of amino acids displayed increased levels in Col-0 at 24 hours p.i. (Supplementary Table S8). In contrast, tryptophan showed no significant induction in *med16* and deficient induction in *med18*. The lack of tryptophan-induction in *med16* was mainly caused by elevated levels already before infection (Fig. 5A). In contrast, *med18* was unable to induce tryptophan levels in response to infection. Furthermore, the elevated leucine levels observed in Col-0 was missing in *med16*, *med18* and *cdk8*. Finally, the fold induction of phenylalanine was similar in Col-0 and mutants, but its absolute levels were elevated in *med16* relative to the other strains in both infected and mock-infected plants (Fig. 5B).

Four benzenoids were induced in Col-0 at 24 hours p.i. relative to the mock-infected control (Supplementary Table S8). One was SA, which also showed induced levels in all mutants. However, we detected increased absolute levels of SA in *med18*, *med25* and *cdk8* relative to Col-0 in mock-infected plants (Fig. 5C). SA is produced by hydroxylation at the second position of the benzene ring of benzoic acid and can thus also be named 2-OH-benzoic acid. Additional hydroxylation at one or several other positions on the benzene ring creates different benzenoids such as 4-OH-bensoic acid and 2,4-di-OH-benzoic acid. Conjugation of the hydroxylated benzoic ring to a glucose moiety results in formation of the hydroxy-benzoic acid glucosides SAG (Salicylic acid glucoside) and SGE (Salicylic acid glucose ester). Such modifications have been suggested to affect translocation into different cellular compartments (Maruri-López et al., 2019). We identified two different glycosylated forms of SA (Salicylic acid glycoside_1 (SAG_1) and Salicylic acid glycoside_2 (SAG_2) in our LCMS assays. (Supplementary Fig. S3A-S3B). SAG_1 was present at low levels in Col-0 and did not increase upon infection while SAG_2 was highly induced. Interestingly, SAG_2 levels were increased in *med25.* Even though Col-0 showed a higher fold-induction of SAG_2 (3.3-fold) compared to *med25* (2.0-fold) at 24 hours p.i., the absolute levels were the same in mock-infected *med25* as in infected Col-0 (Supplementary Fig. S3B). Similar elevated levels in mock-infected *med25* were also found for three other metabolites that represent modified versions of benzoic acid: 4-hydroxybenzoic acid, 2,4-dihydroxybenzoic acid and Protocatechuic acid 3-glucoside (Supplementary Fig. S3C-S3E). This suggests that elevated levels of SA-related metabolites in *med25* before infection might contribute to its resistance against *P. syringae* infections.

ICS1 is the key enzyme for pathogen-induced SA production (see Supplementary Fig. S4A for illustration of the SA signaling pathway). Using qPCR, we found that the fold induction of *ICS1* mRNA was significantly higher in *med25* relative to the other lines at 24 hours p.i. (Supplementary Fig. S4B). Furthermore, the expression level of *EDS1* mRNA, which encodes one of the transcriptional activators of *ICS1* expression, was increased in *med25* relative to Col-0, *med16* and *med18* at 24 h p.i., while it was reduced in *cdk8* (Supplementary Fig. S4C). Interestingly, EDS1 was recently shown to physically interact with the Mediator kinase module subunit CDK8 (Chen et al., 2021). In contrast, the other *ICS1* transcriptional activator, *PAD4* did not show significant differences in any mutant relative to Col-0. Responses downstream of SA are mediated by NPR1, a transcriptional co-activator for SA-dependent genes like *NIMIN1* and *PR1*. We found that transcript levels of *NIMIN1* and *PR1* at 24 hours p.i. was similar in *med25* and Col-0 but impaired in *med16*, *med18* and *cdk8* (Supplementary Fig. S4D).

Levels of oxylipins with conjugated OPDA (e.g. MGMG(18:4;O/0:0) / MGMG(12-OPDA)_1 and DGMG(18:4;O/0:0) / DGMG(12-OPDA)_1) were induced at 24 hours p.i. in Col-0, *med16* and *cdk8* (Fig. 5D, Supplementary Table S8). In contrast, they were uninduced in *med18* and *med25*, albeit by different mechanisms. In mock-infected plants, levels were similar in *med18* and Col-0, but they were not induced in *med18* at 24 hours p.i. Mock-infected *med25* on the other hand, displayed levels equivalent to those detected in Col-0 at 24 hours p.i. but showed no increase in response to infection. As for the SA-related metabolites, increased levels of OPDA-conjugated oxylipins in uninfected *med25* might contribute to its resistance to *P. syringae* infection.

Camalexin and a set of modified, related metabolites were among the most highly induced in Col-0 at 24 hours p.i. (Fig. 5E, Supplementary Table S8). However, Camalexin induction was impaired in all mutants, especially *med25* and *cdk8*. Again, we found that these differences were due to different mechanisms. In mock-infected *med25*, Camalexin was elevated to even higher levels than those observed in Col-0 after infection. In contrast, *cdk8* displayed low levels in mock-infected plants and no increase at 24 hours p.i. This might contribute to their opposite phenotypes. The difference in Camalexin levels between *med25* and *cdk8* is likely due to defects in expression of *PAD3*, which encodes the key enzyme in the Camalexin synthesis pathway (Supplementary Fig. S5A). *PAD3* expression was higher in uninfected *med25* but lower in uninfected *cdk8* relative to Col-0 (Supplementary Fig. S5B). Finally, we observed induction of some phytoprostanes and AMP in Col-0 at 24 hours p.i. Their induction was slightly defect in *med16* and *med18* (Supplementary Table S8).

### Sugars, amino acids and jasmonates accumulate in *med16* and *cdk8* but decrease in *med25* at the late stage of infection

We made scatterplots representing changes in levels of all 222 identified metabolites at 72 hours p.i. relative to control. The distribution of log_2_-FCs was compared between each mutant and Col-0 using simple linear regression (Fig. 6A). To reveal deviations in mutants, we tested the null hypothesis that the slope of the linear regression was equal to one. Table 1 shows slope estimates and adjusted R∧2 for the regression as well as the p-value for slope equal one. *med16* and *cdk8* had a significantly lower slope of 0.826 and 0.720 indicating that their metabolic responses were more affected by infection compared to Col-0. In contrast, *med25* had a slope of 1.819 indicating that it was less affected than Col-0. The slope for *med18* was not significantly different from slope equal one, indicating that it behaves as Col-0. These results correlate with the phenotypic differences of these mutants relative to Col-0 in response to *P. syringae* infection.

**Figure 6.**
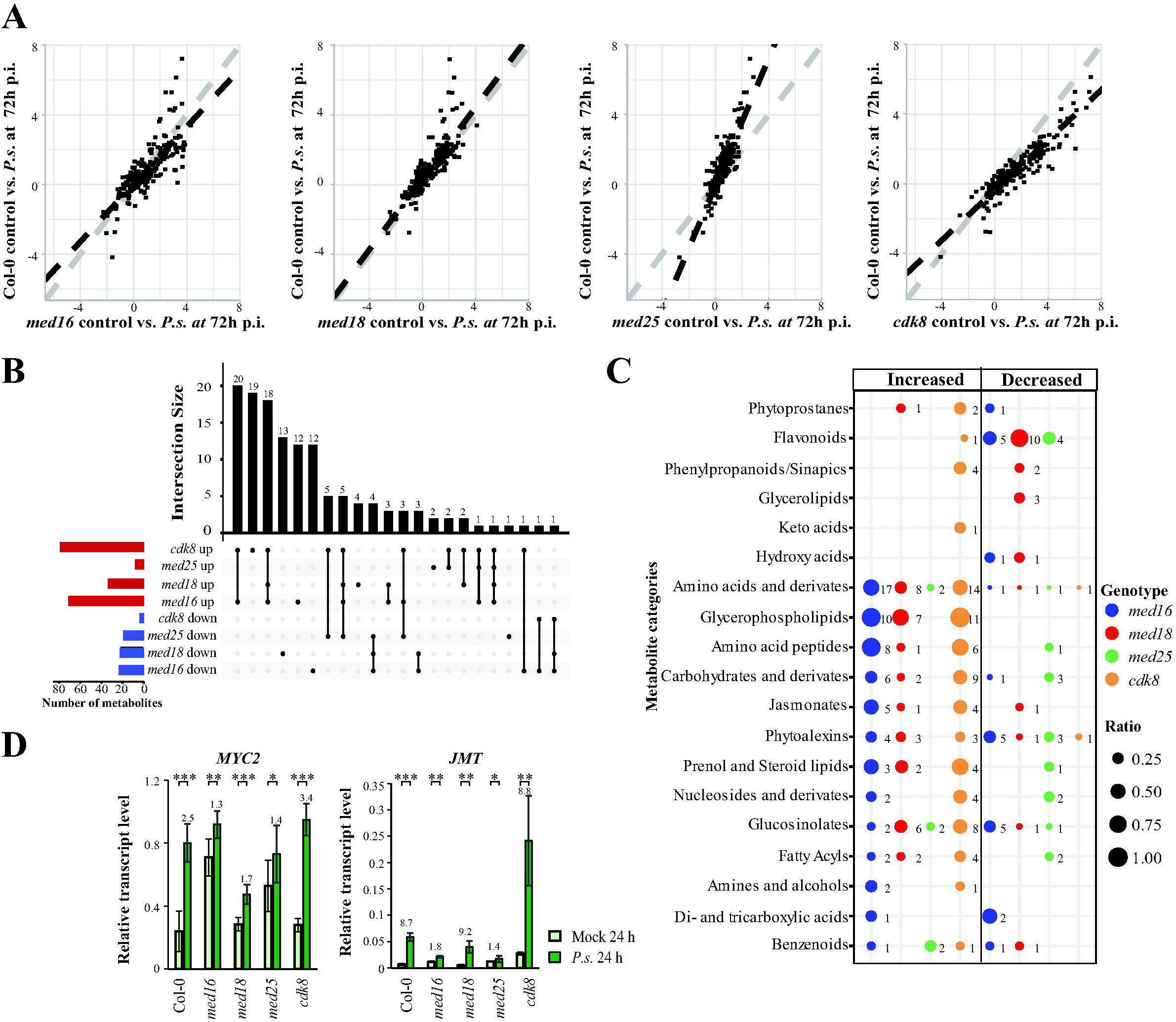
*med25* shows an attenuated infection response whereas *med16* and *cdk8* show the highest alterations in metabolite levels compared to Col-0 at the late stage of infection. (**A**) Scatter plots of the relationship between metabolite changes at 72 hours p.i. (relative to control at 72 h) between each mutant (x-axis) and Col-0 (y-axis). (**B**) Upset diagram illustrating the overlap and uniqueness in increased and decreased metabolite levels in each mutant relative to Col-0 at 72 hours p.i. Total number of metabolites displaying changed levels in each mutant are represented as red (increased) and blue (decreased) bars to the left. (**C**) Distribution of metabolites displaying changed levels in each mutant for each of the nineteen metabolite categories. Sizes of circles represent the ratio of the of the number of metabolites in each category that display significantly distinct levels in mutants relative to Col-0 and the total amount of metabolites detected for each category in our full data set of 222 metabolites. Numbers to the right of each circle represent the total number of metabolites that show significantly changed levels in each mutant. In (**A**) a significance cutoff of log_2_-FC > 0.5 or<-0.5 and p < 0.05) was used. In (**B**) and (**C**) a significance cutoff of log_2_-FC > 1 or <-1 and p < 0.05) was used. (**D**) Levels of two JA activated genes (*MYC2* and *JMT*) in mock-infected and infected cells at the 24-hour time point. Asterisks indicate significant differences (*: p< 0.05, **: p< 0.01, ***: p< 0.001) and the error bars show the SD of three biological replicates.

**Table 1.**
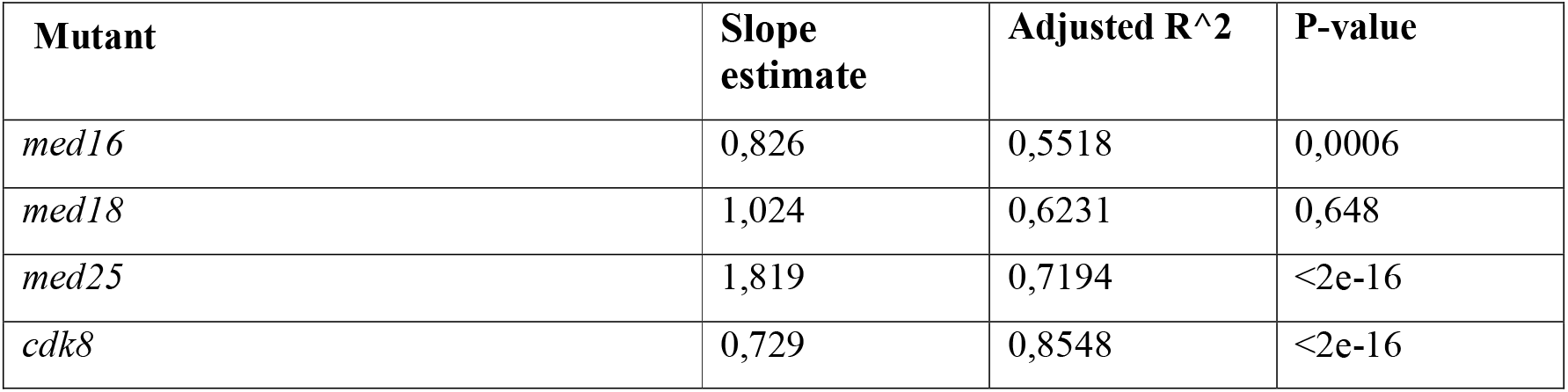
Summary of linear regression estimates of log_2_-fold changes for mutants relative to Col-0.

| **Mutant** | **Slope estimate** | **Adjusted R^2** | **P-value** |
| --- | --- | --- | --- |
| *med16* | 0,826 | 0,5518 | 0,0006 |
| *med18* | 1,024 | 0,6231 | 0,648 |
| *med25* | 1,819 | 0,7194 | <2e-16 |
| *cdk8* | 0,729 | 0,8548 | <2e-16 |

For metabolites that were increased in each mutant relative to Col-0 at 72 hours p.i., we found a large overlap of 48 metabolites between *med16* and *cdk8* (Fig. 6B). Twenty-four of them were also increased in *med18* (Supplementary Table S9). In line with its opposite phenotype, only three were increased while eight were reduced in *med25*. Using a less strict threshold we identified a set of amino acids which were increased in *med16* and *cdk8* but decreased in *med25* (Supplementary Fig. S6A). The fold change for aspartic acid showed the reverse, being reduced in *med16*, and *cdk8* but increased in *med25* compared to Col-0. We also identified twenty-five lipids that where commonly increased at the 72-hour time point in *med16*, and *cdk8* and one of them, the unsaturated lipid stigmasterol, was reduced in *med25.* Thus, stigmasterol levels correlate to bacterial growth in each mutant relative to Col-0. Finally, a set of twelve carbohydrates showed an increased fold-change in *med16* and/or *cdk8*, and half of them were reduced in *med25*. In particular, trehalose had the highest log_2_-fold increase compared to Col-0 in *cdk8* (log_2_FC= 3.7) but was decreased in *med25* (log_2_FC= -1.4).

The sensitive *med16* and *cdk8* mutants displayed increased accumulation of jasmonates after infection (Fig. 6C). *med16* showed a higher fold accumulation of 12-hydroxylated-JA-Ile, and *cdk8* displayed a pronounced increase of methylated JA (MEJA), while both mutants showed induced levels of JA-Ile (Supplementary Fig. S6B). α-linolenic acid (18:3), a fatty acid substrate of jasmonate biosynthesis, was also elevated in the *med16* and *cdk8* compared to Col-0 (Supplementary Fig. S6B), suggesting a broad and dysfunctional regulation of the jasmonate biosynthetic pathway in these mutants. The increased levels of jasmonates in *med16* corroborated the increased mRNA levels of JA-related genes in *med16* relative to the other mutants already in mock-infected plants (Supplementary Fig. S6C).

Many JA-induced genes are regulated by MYC2. Interestingly, we found that the levels of *MYC2* mRNA in both *med16* and *med25* under control conditions were elevated to those we detected in Col-0 at 24 hours p.i. However, the fold induction of *MYC2* was reduced in *med16* and *med25*, resulting in similar levels of *MYC2* in Col-0, *med16*, *med25* and *cdk8* at 24 hours p.i. (Fig. 6D). MED25 has been reported as important for MYC2-dependent gene regulation (Chen et al., 2012). Accordingly, induction of *JMT*, which is a MYC2 target gene encoding an S-adenosyl-L-methionine:jasmonic acid carboxyl methyltransferase that catalyzes the formation of MEJA from JA, was deficient in *med16* and *med25* due to a combination of elevated levels before infection, and deficient induction after infection (Fig. 6D). *cdk8* displayed elevated *JMT* mRNA levels before infection but in contrast to *med16* and *med25*, it showed the same fold-induction as Col-0. Thus, elevated MEJA levels could result from increased *JMT* expression in *cdk8*. Finally, two other MYC2 target genes, *JAZ6* and *LOX2*, were uninduced in *med25* after infection (Supplementary Fig. S6D). The same impairment was detected for *med16* after infection whereas *cdk8* showed an accumulation of *JAZ6*.

## DISCUSSION

Hostile environmental conditions result in substantial losses to forestry and food production worldwide. Increased temperature and reduced water availability are important abiotic factors, while biotrophic bacteria and necrotrophic fungi are central biotic factors limiting plant growth and productivity. Due to their sessile nature and lack of specialized immune cells, plants have developed sophisticated ways to alter their metabolism to adapt to environmental changes. Following perception of stress by different receptors, signals are relayed to the nucleus via complex cellular signaling networks. This involves second messengers such as reactive oxygen intermediates (ROIs), calcium-associated proteins and mitogen-activated protein (MAP) kinase cascades and leads to induction of stress tolerance genes and changes in protein and metabolite levels.

Mediator plays important roles for responses to biotic and abiotic stresses. Mutations in *MED2*, *MED14*, *MED16*, *MED18*, and *MED25* causes impaired drought, high salinity and cold responses (Bäckström et al., 2007; Hemsley et al., 2014), and MED8, MED14, MED15, MED16, MED18, MED20, and MED25 are involved in biotic stress responses (Kidd et al., 2009; Elfving et al., 2011; Fallath et al., 2017). Furthermore, Mediator subunits are important for the Salicylic acid (SA) and Jasmonic acid/Ethylene (JA/ET) signaling pathways, which control several aspects of defense signaling in plants (Zhang et al., 2012; Caillaud et al., 2013; Dhawan et al., 2009). Interactions between specific Mediator subunits and regulatory TFs are important for responses to biotrophic pathogens such as *P. syringae* (Seo et al., 2019; Huang et al., 2019). Less is known regarding how Mediator mutations affect metabolite levels and how such changes influence infection susceptibility. Our results show that *med16* and *cdk8* are susceptible to *P. syringae* infection while *med25* shows increased resistance compared to Col-0 plants. We identify specific metabolites that can explain the phenotypic variations and show that differences in metabolite responses between Col-0 and mutants can result from pre-existing differences already in uninfected plants, or in how they change in response to infection.

In uninfected cells, three metabolite categories showed differences in mutants relative to Col-0 and thus might explain the variations in phenotypes for each of *med16*, *med25* and *cdk8*. Phytoalexins, carbohydrates and GLSs were reduced in the susceptible *med16* and *cdk8*, while they were increased in the more resistant *med25*. This agrees with previous reports showing that the levels of these classes of metabolites are induced in Col-0 in response to infections (Tsuji et al., 1992; Tierens et al., 2001; Trouvelot et al., 2014). In particular, GLSs are known to increase in infected Col-0 and to play important roles in infection defense (Andersson et al., 2014). We found reduced levels of Sulforaphanes in uninfected, susceptible *med16* and *cdk8* but increased levels in the more resistant *med25*. Sulforaphane is produced in response to several non-host bacterial pathogens, and *P. syringae* strains adapted to Arabidopsis express *SURVIVAL IN ARABIDOPSIS EXTRACTS* (*SAX*) genes, which enable them to detoxify host-produced sulforaphane. In addition, we identified two more GLSs (Hirsutin and Neoglucobrassicin) that showed reduced levels in uninfected *med16* and *cdk8* and additional GLSs, carbohydrates and phytoalexins that were uniquely reduced in either *med16* or *cdk8*. In line with our metabolite profiling of uninfected Col-0 and mutants, our RNA-seq analyses showed that *med16* and *cdk8* have reduced levels of transcripts encoding enzymes involved in GLS biosynthesis and metabolism and expression of *PAD3* which encodes a key enzyme in synthesis of Camalexin was higher in uninfected *med25* but lower in *cdk8* relative to Col-0. Finally, a set of metabolites showed elevated levels uniquely in uninfected *med25*. The most common categories were benzenoids, which are precursors for SA synthesis. We conclude that levels of GLSs and phytoalexins in mock-infected mutants correlate with their *P. syringae* infection susceptibility. Since SA plays play a major role in defense against *P. syringae* (Vlot et al., 2009), the elevated levels of several SA-related metabolites in *med25* before infection might contribute to its resistance against *P. syringae* infections.

In the early response to *P. syringae,* we identified a further set of metabolites whose changes in levels deviate in mutants relative to Col-0. One important category was amino acids, which play important functions as precursors for several defense-mediating secondary metabolites. Tryptophan is an important precursor for synthesis of e.g. Camalexin, indolic GLSs, and auxin (indole-3-acetic acid; IAA). Our results show that tryptophan levels are increased in *med16* already before infection to the same levels as those we observe in Col-0 and the other mutants after infection. Phenylalanine, a precursor for e.g. flavonoids, anthocyanins, benzoic acids and SA, was induced to the same levels in Col-0 and all mutants but its absolute levels were increased in *med16* both before, and at the early timepoint after infection. A set of benzoic acids, among them SA (or 2-OH-benzoic acid), showed dysregulation in the mutants at the early timepoint after infection. We identified two different glycosylated forms of SA (SAG_1 and SAG_2) in our LCMS assays which were differently regulated. SAG_1 was expressed at low levels in both Col-0 and *med25*, while SAG_2 was similarly induced in Col-0 and *med25* but present in much higher levels in the mutant. These metabolites likely correspond to Salicylic acid glucoside (SAG) and Salicylic acid glucose ester (SGE). Based on previously reported results showing that levels of SGE in Arabidopsis is lower than SAG and that SGE is much less induced in response to wounding compared to SAG (Ogawa et al., 2010), it is likely that SGE in our assays corresponds to SAG_1 and SAG to SAG_2. Camalexin, which we identified as increased in uninfected *med25* but reduced in *cdk8* relative to Col-0, was also impaired in its induction at the early timepoint after infection, especially in *med25* and *cdk8,* and we obtained similar results for set of modified Camalexin-related metabolites. Again, the mechanisms for the lack of induction differed between mutants. Uninfected *med25* displayed Camalexin levels even higher than infected Col-0 and they did not increase further upon infection. In contrast, *cdk8* showed low Camalexin levels both before and after infection. Finally, oxylipins with conjugated OPDA were induced in Col-0, *med16* and *cdk8* at the early time point after infection but did not increase in *med18* and *med25*.

At the late time point after infection, we observed several differences. Generally, *med16* and *cdk8* showed larger effects on metabolite levels after infection relative to Col-0, while *med25* showed less. Nearly fifty metabolites were present at higher levels at the late time point in both *med16* and *cdk8* relative to Col-0, while they were unaffected or present at lower levels in *med25*. Of these, we again identified a set of amino acids as well as twenty-five lipids that were commonly increased at the 72-hour timepoint in *med16*, and *cdk8* relative to Col-0. One of them, the unsaturated lipid stigmasterol, was also reduced in *med25.* Stigmasterol is synthesized at pathogen inoculation sites where it integrates into plant cell membranes and favors susceptibility to bacterial pathogens (Griebel and Zeier, 2010). Also, a set of carbohydrates showed increased levels in infected *med16* and *cdk8*, while they were decreased in *med25*. In particular, trehalose levels were highly increased in *cdk8* but severely decreased in *med25*. Application of exogenous trehalose was recently shown to increase susceptibility to *P. syringae* (Wang et al., 2019), suggesting that its elevated levels at 72 hours p.i. might contribute to the increased and decreased bacterial content observed in *cdk8* and *med25*, respectively. Finally, we identified increased levels of jasmonates in *med16* and *cdk8* relative to Col-0 and the other mutants at the late time point after infection. Jasmonates comprise a family of oxylipins including JA and a set of derivatives that regulate various aspects of plant immunity and development. It is therefore likely that the dysregulation of JA we observe contribute to their phenotypic behavior in response to infection.

Here we used a combination of metabolomics and RNA-seq and four Mediator mutants to study molecular mechanisms for how Arabidopsis respond to infection caused by the biotrophic pathogen *P. syringae*. We identify a set of unique metabolites and metabolite categories that show different types of defects in mutants relative to wild type Arabidopsis and we show that these differences are often based on defects in regulation or function of genes encoding key enzymes or regulatory TFs that control metabolic pathways which are important for a proper infection response. Our results form a base for future research that will hopefully result in plants and crops which are more resistant to infections.

## ACKNOWLEDGEMENTS

The Metabolic profiling was performed at the Swedish Metabolomics Center in Umeå, Sweden (https://www.umu.se/en/research/infrastructure/metabolomics/).

## AUTHOR CONTRIBUTIONS

J.B, Å.S, M.R and S.B planned and designed the research. J.B, F.K and Q.M.I performed experiments. J.B, V.J, A.V, Å.S, M.R. and S.B analyzed data. J.B and S.B wrote the manuscript with the assistance of all co-authors.

## DATA AVAILABILITY

The sequencing data for the *med16*, *med18*, *cdk8* mutants and the wild type Col-0 has been deposited at the European Nucleotide Archive (ENA, www.ebi.ac.uk/ena) under accession number PRJEB33339. The sequencing data for the *med25* mutant and the wild type Col-0 has been deposited at https://figshare.com/s/9a58efd7fee99b1d755c. Similarly, The Metabolomics data have been deposited to the EMBL-EBI MetaboLights database (DOI: 10.1093/nar/gks1004. PubMed PMID: 23109552) with the identifier MTBLS8105. The complete dataset can be accessed here: http://www.ebi.ac.uk/metabolights/MTBLS450.MTBLS8105.

